# Biochemical Basis for the Regulation of Biosynthesis of Antiparasitics by Bacterial Hormones

**DOI:** 10.1101/2020.04.27.064931

**Authors:** Iti Kapoor, Philip Olivares, Satish K. Nair

**Affiliations:** Department of Biochemistry, University of Illinois at Urbana Champaign, 600 S. Mathews Ave, Urbana IL USA; Institute for Genomic Biology and University of Illinois at Urbana Champaign, 600 S. Mathews Ave, Urbana IL USA; Center for Biophysics and Computational Biology, University of Illinois at Urbana Champaign, 600 S. Mathews Ave, Urbana IL USA

## Abstract

Diffusible small molecule microbial hormones drastically alter the expression profiles of antibiotics and other drugs in actinobacteria. For example, avenolide (a butenolide) regulates production of avermectin, derivatives of which are used in the treatment of river blindness and other parasitic diseases. Butenolides and γ-butyrolactones control production of pharmaceutically important secondary metabolites by binding to TetR family transcriptional repressors. Here, we describe a concise, 22-step synthetic strategy for the production of avenolide. We present crystal structures of the butenolide receptor AvaR1 in isolation, and in complex with avenolide, as well as AvaR1 bound to an oligonucleotide derived from its operator. Biochemical studies guided by the co-crystal structures enable identification of 90 new actinobacteria that may be regulated by butenolides, two of which are experimentally verified. These studies provide a foundation for understanding regulation of microbial secondary metabolite production, which may be exploited for the discovery and production of novel medicines.

## INTRODUCTION

The rise of drug-resistant pathogens continues to compromise human health and is exacerbated by the decline in the discovery rate for new anti-infectives.^1,2^ A major limitation is the lack of tools that enable access to the vast library of bacterial natural product antibiotics. Despite the fact that statistical surveys depict the number of antibiotics that are genetically encoded within the *Streptomyces* genus to be in excess of ~300,000 new molecules, a large repertoire of these compounds cannot be produced when the strain is grown under standard laboratory conditions.^3,4^ The responsible biosynthetic genes are ‘silent’ under laboratory condition and are regulated via unknown mechanisms.^5,6^

The diffusible small molecule γ-butyrolactone (GBL) A-factor plays an essential role in the biosynthesis of the antibiotic streptomycin in *Streptomyces gresius*. The intracellular target of A-factor has been identified as a member of the TetR family, and these receptors are shown to regulate of antibiotic biosynthesis in in several actinobacterial species (Figure 1A).^6–9^ The use of exogenous GBLs has been shown to induce secondary metabolite production from otherwise silent clusters.^10,11^ However, the alkali labile nature of γ-butyrolactones, and the pleiotropic nature of GBL-mediated regulation limit the general use of these hormones.

**Fig. 1.**
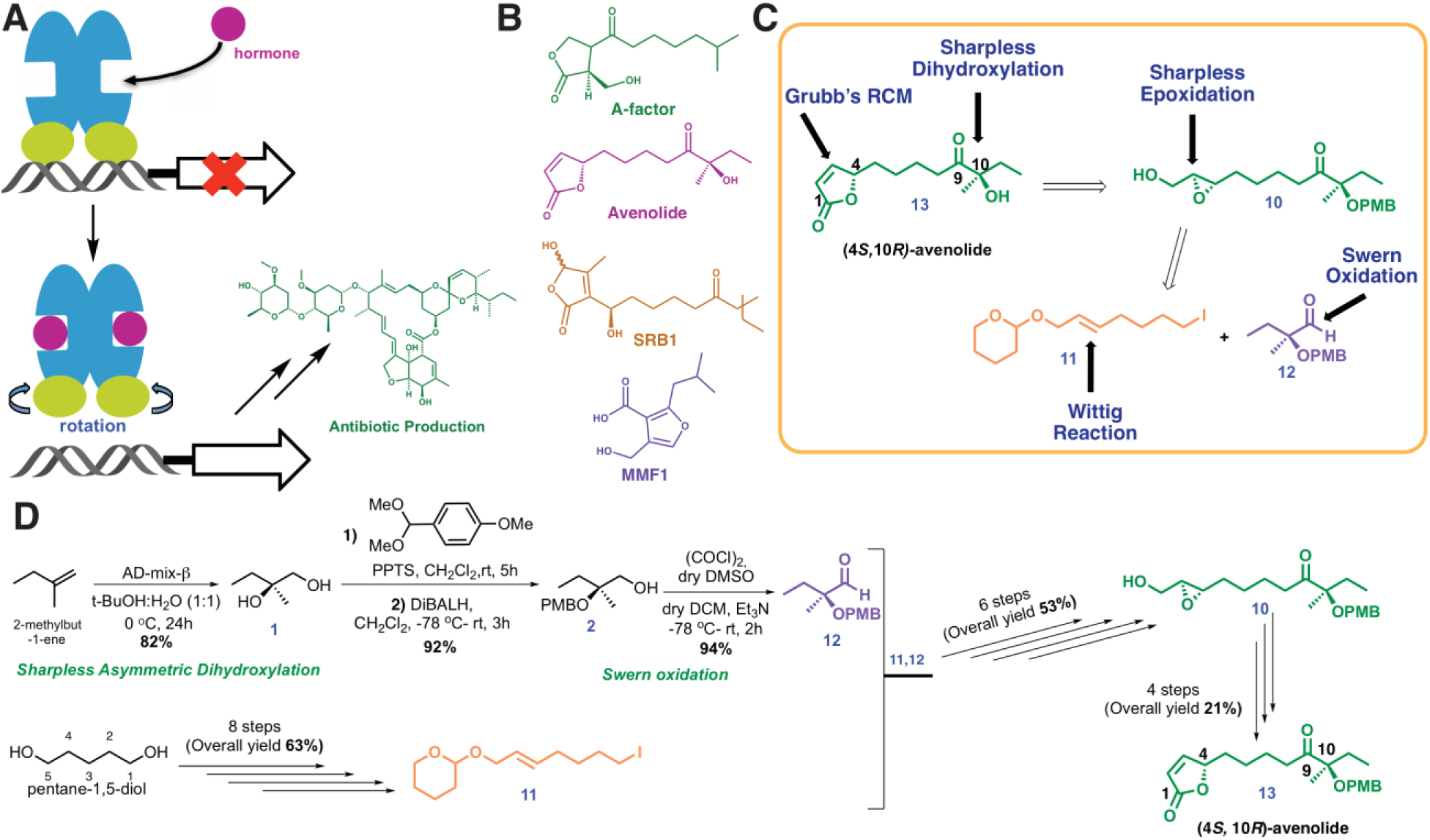
Chemical structures and retrosynthetic scheme for avenolide. **A**. Representation of the mechanism for hormone induced transcriptional activation in bacteria. A-factor is a γ-butyrolactone, avenolide is an alkylbutenolide, SRB1 is a 2-alkyl-3-methyl-4-hydroxybutenolide, and MMF1 is a 2-alkyl-4-hydroxymethylfuran-3-carboxylic acid. **B.** Structures of representative compounds from the four known classes of bacterial hormones. **C**. Retrosynthetic scheme for avenolide synthesis involving five key reactions. **D**. Overall summarized and synthetic scheme for total synthesis of (4*S*,10*R*)-avenolide with total number of steps and reaction yields.

Related class of bacterial hormones include the 2-alkyl-3-methyl-4-hydroxybutenolides, the 2-alkyl-4-hydroxymethylfuran-3-carboxylic acid, and alkylbutenolides (Figure 1B). Butenolides, such as avenolide (Figure 1B), triggers the production of secondary metabolites with a minimum effective concentration in the low nanomolar range in *S. avermitilis*.^12,13^ Notably, butenolides show greater pH stability and generally regulate fewer processes as compared to γ-butyrolactones. The recent discovery of avenolide activity observed in about 24% of actinomycetes (n=51), suggests that other active actinobacteria can also produce avenolide-like compounds to regulate secondary metabolism.^10^ For example, avenolide regulates the production of the anthelminitic compound ivermectin.^14,15^ Ivermectin is on the World Health Organization’s list of essential medicines, and has lowered the incidence of otherwise untreatable parasitic infections, including river blindness, strongyloidiasis, and lymphatic filariasis (elephantiasis).^16,17^

While enzymatic and synthetic routes towards the production of γ-butyrolactones have been described, access to butenolides has been restrictive.^18,19^ Here, we describe a concise 22-step convergent route towards the total synthesis of avenolide, enabling biochemical and biophysical characterization of its interaction with the AvaR1 receptor. We also present structures of AvaR1, in isolation (2.4 Å resolution), in complex with avenolide (2.0 Å resolution), and bound to a synthetic DNA oligonucleotide (3.09 Å resolution) derived from its natural binding site. Using the primary sequence of AvaR1 and synteny of genes that are likely involved in butenolide biosynthesis, we identify 89 additional putative butenolide receptors. Mapping of residues at the ligand-binding site with sequence conservation highlights their importance in ligand activation. The identification of these putative avenolide-responsive strains may enable the production of novel metabolites in the presence of the hormone. As proof of principle, we show that the supplementation of synthetic avenolide into growing cultures of two strains that contain homologous receptors results in visible changes to the production media.

## RESULTS AND DISCUSSION

### Convergent Synthesis of (4*S*, 10*R*)-Avenolide and Characterization of Binding to AvaR1

A retro-synthetic strategy was pursued to allow for the production of avenolide from the convergent synthesis of three key fragments consisting of the iodide **11**, the aldehyde **12** and epoxy **10** (Figure 1C) following the reported protocol by Uchida *et al.*^20^ Starting from the commercially available 1-methyl-2-butene, a Sharpless asymmetric dihydroxylation^21^ produced the diol intermediate **1** in 82% yield in a single step (Figure 1D). The desired key intermediates aldehyde **12** and iodo alkene **11** and epoxy **10** were also made following the same reported protocol except the use of TBDPS rather than TBS for improved reaction monitoring using TLC. **10** was then stereospecifically converted into allyl alcohol **17** in a single step, by using titanocene dichloride and Zn powder.^22,23^ The final product, stereospecific (4*S*, 10*R*)-avenolide, **13** was then produced by ring-closing metathesis of **19** treating the dialkene with Grubb’s second-generation catalyst.^24^ The final yield of avenolide was 14 mg total from 15 g of starting material. The identity of all intermediates, as well as that of the final product, was determined using ^1^H-NMR and ^13^C NMR which matched with the reported data. Detailed experimental methods and NMR spectra can be found in supporting information.

### Crystal structure of AvaR1 and the binary complex with avenolide

The structure of AvaR1 was determined to 2.4 Å resolution (Figure 2A) using crystallographic phases determined from anomalous diffraction data collected from SeMet-labeled protein crystals. The overall structure is reminiscent of that of other TetR-family transcriptional repressors, and consists of an obligate homodimer.^25^ Each monomer is entirely helical and consists of a DNA-binding domain (composed of α helices 1-4), and a ligand-binding domain (consisting of α helices 5-13). The dimer interface is formed via interactions between the two ligand-binding domains and is formed mainly through hydrophobic packing interactions.

**Fig. 2.**
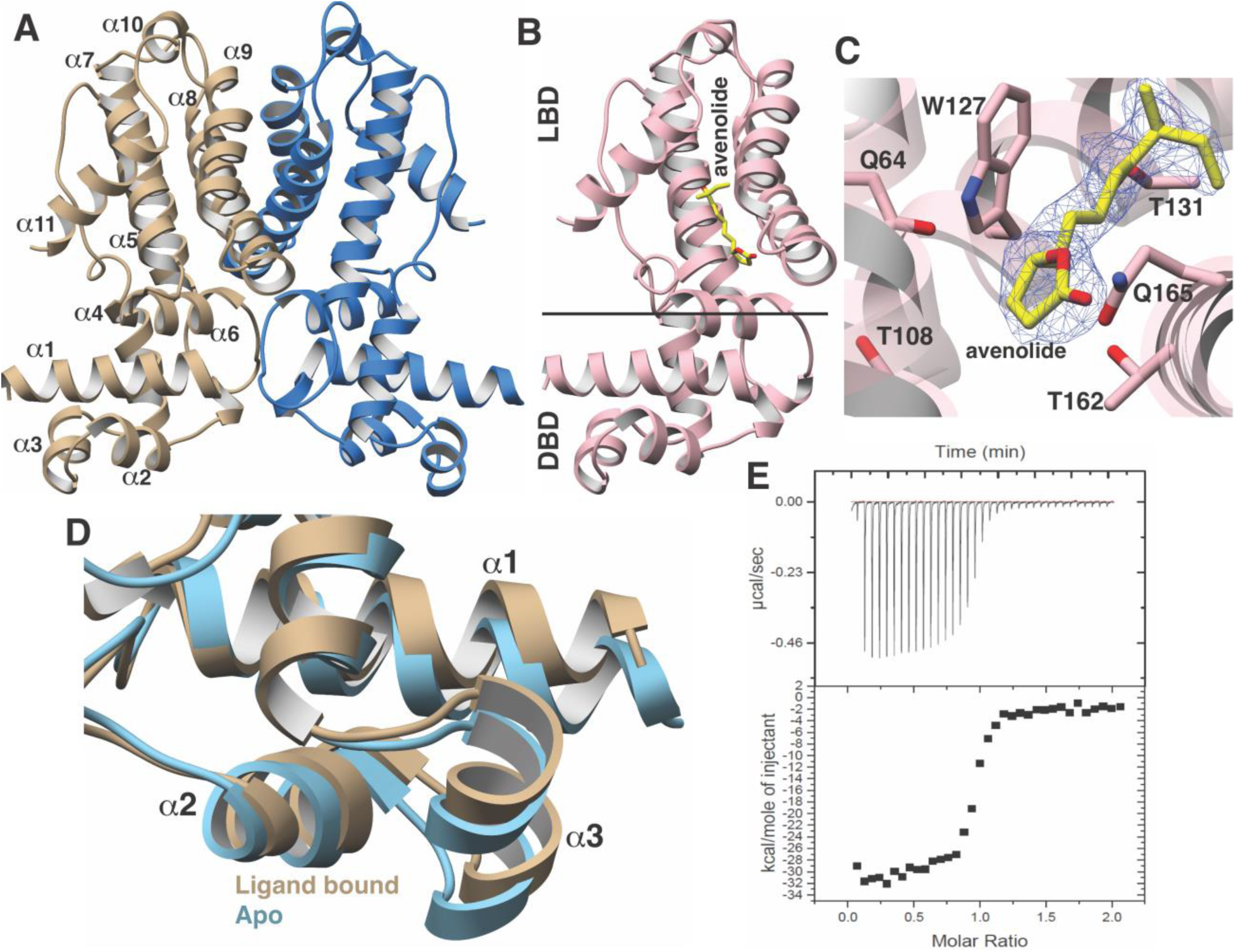
Structural characterization of AvaR1-avenoide binding interaction. **A**. Structure of the AvaR1 homodimer in the absence of bound ligand. One monomer shown colored in blue and another in tan. **B**. Co-crystal structure of one monomer of AvaR1 (in pink) bound to (4*S*,10*R*)-avenolide (in yellow ball-and-stick). The ligand binding domain (LBD) and the DNA binding domain (DBD) are indicated. **C**. Difference Fourier map (countered at 3) calculated with coefficients |F(obs)|-|F(calc)| with the coordinates of the avenolide omitted prior to one round of refinement. The coordinates of the final structure are superimposed. **D**. Superposition of the structures of the DBD of AvaR1 in the presence (tan) and absence (cyan) of bound ligand. Ligand binding induces a 10° shift in this domain that would preclude DNA binding. **E**. Representative binding isotherm for the interaction of AvaR1 with (4*S*,10*R*)-avenolide indicative of a 1:1 binding stoichiometry.

Cocrystallization efforts of AvaR1 bound to avenolide yielded crystals that diffracted to 2.0 Å resolution, and crystallographic phases were determined using molecular replacement (Figure 2B).^26^ Clear density for the entire hormone can be visualized bound to the ligand-binding domain of both monomers in the homodimer (Figure 2C). Notably, the structure of AvaR1 has undergone conformational shifting upon ligand binding, and a structure-based superposition against the ligand free structure illustrates that binding of the hormone results in a ~10° shift in the DNA-binding domains of each monomer (Figure 2D). This shift results in an increase in the distance between the two DNA binding domains in the dimer upon binding of the ligand, which would preclude DNA binding by the ligand-bound homodimer. Additional local changes between the two structures included the movement of Gln165, which swings into the binding pocket to make hydrogen-bonding interactions with the lactone ring, as well with Gln64. Lastly, Thr108, which is harbored on helix 6, also moves to accommodate interactions with the hormone. The indole side chain of Trp127 likewise shifts to increase the volume of the binding cavity. Other interactions include hydrogen bonds between Thr131 and the C10 hydroxy, and Thr162 that interacts with the lactone ring of avenolide. The superposition suggests that the new contacts formed by these residues likely couple hormone binding to the conformational shift between the two monomers of AvaR1 (Figure 2D).

We used isothermal titration calorimetry to characterize the binding interaction between the synthetic avenolide and AvaR1 (measurements were conducted in triplicate). The resultant binding isotherms shows the point of inflection at a molar ratio of N-1, which suggests a 1:1 binding of ligand per monomer (Figure 2D). The strength of the binding is measured to be K_d_ = 42.5 nM ± 2.1 nM (3 independent trials), which correlates with the reported value of ~4 nM that was estimated from gel shift-based assays.^14^

### Structure based comparison across different GBL-like receptors

A structure-based sequence alignment of GBL-like receptors for which ligand specificity has been established reveals a strong conservation of residues shown to be critical for ligand binding, suggesting a common mechanism for ligand recognition across disparate classes of receptors (Figure 3A). Residues that surround the alkyl chain of avenolide include Trp127, Val158, and Phe161, which are almost universally conserved across all members of the receptor family, while Leu88, which forms the opposite wall of the binding cavity is always a hydrophobic residue but of variable size (Figure 3B). The chain length of the methylenomycin furans (MMFs) is shorter than those of the GBLs and butenolides; correspondingly, residues at the base of the ligand-binding cavity in the AvaR1, such as Ala85, His130, and Thr131, are replaced by bulkier Glu107, Leu150, and Leu151, in the sequence of the MMF receptor MmrF. Residue Thr161 is within hydrogen bonding distance to the lactone ring of avenolide, and this residue is conserved in receptors that bind to γ-butyrolactones and butenolides, but absent in receptors for other classes of hormones such as MmrF. Likewise, Gln64 in AvaR1 is positioned on the opposite site of the lactone and is conserved among receptors that bind to structurally related classes of hormones, but is absent in the structure of MmrF. The latter sequence contains a Tyr85 at a near equivalent position, which may be necessary for interactions with the carboxyl group of MMF.

**Fig. 3.**
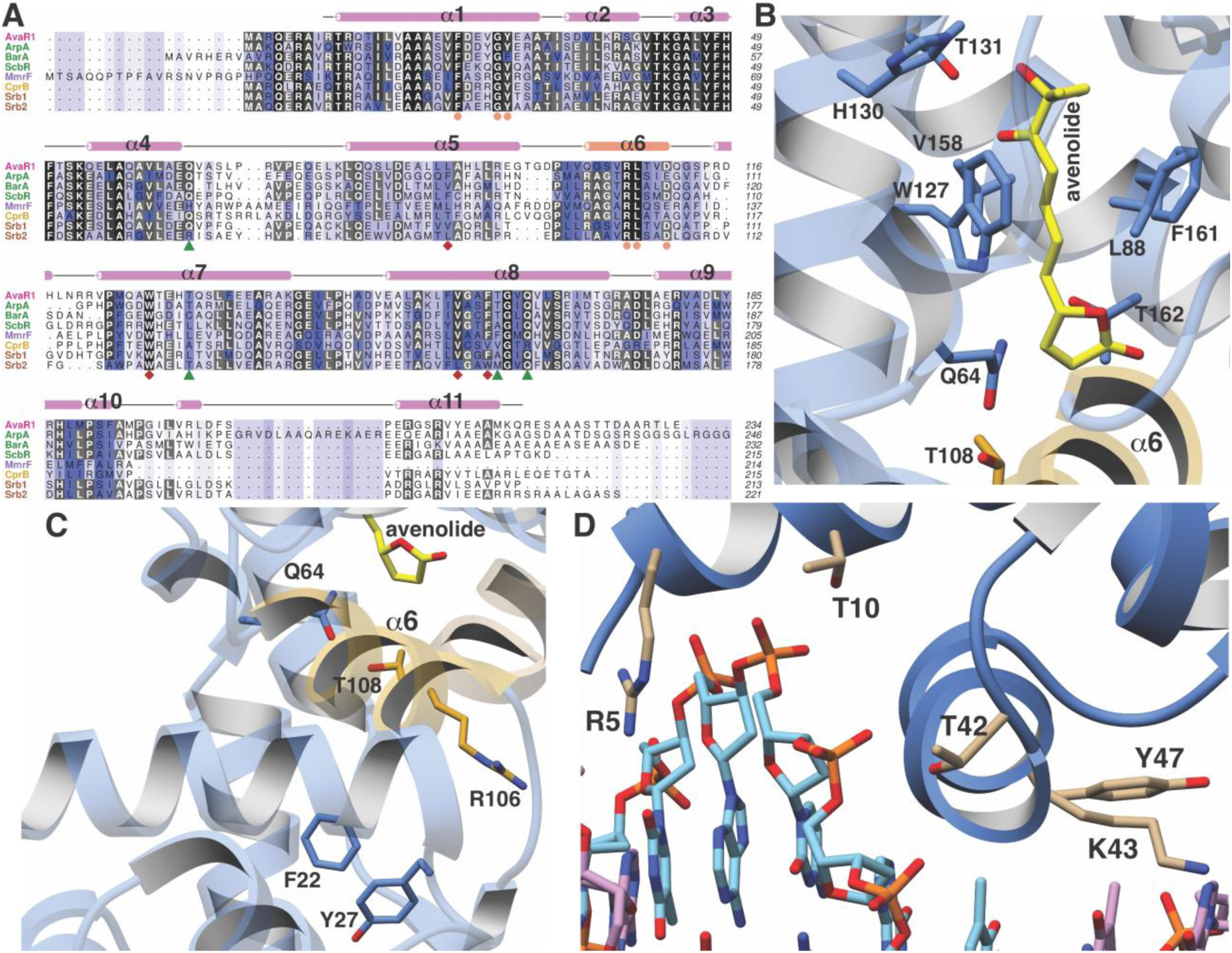
**A.** Multiple sequence alignment of various GBL-like receptors for which ligand specificity is known. The color-coding of the receptor names reflects the ligand class as colored in Figure 1 B. Residues involved in interactions with the lactone are colored in green triangles, those interacting with the alkyl chain are colored as red diamonds, and those proposed to be involved in mediating hormone-dependent conformational movement are shown as orange circles. **B**. Close-up view of the hormone-binding cavity showing residues in contact with the bound ligand. **C**. Spatial orientation of conserved residues that are proposed to induce movement of the DNA binding domain in response to binding of the hormone in the ligand binding domain. **D**. Close-up view of the DNA-binding domain of AvaR1 in complex with the *aco* ARE.

The lactone ring of avenolide is situated above helix α6, which contains residues that are nearly universally conserved among GBL-like receptors, and include Ser103, Val104, Arg105, Leu106, Val107, and Asp108. Notably, this helix bridges the ligand binding domain and the DNA-binding domain, suggesting that is plays a role in coupling ligand binding to DNA dissociation (Figure 3C). Specifically, movement of Gln64 and Thr108 into the ligand-binding cavity of AvaR1 upon engagement of the hormone results in the displacement of helix α6 away from the pocket. The orientation of Arg105, located on the opposite side of helix α6 is established through multiple hydrogen bonding interactions with the backbone carbonyls of conserved residues in helix α1, including the universally conserved Phe22, Gly26, and Tyr27. Hence, accommodation of hormone binding necessitates domain movement of the DNA-binding domain, in order to preserve the suite of hydrogen bonding interactions with helix α6. As noted, Gln64 and Thr107 are largely conserved among receptors that bind lactone-containing hormones, suggesting a common mechanism for coupling ligand binding to DNA dissociation.

### Crystal structure of AvaR1 and the binary complex with the *aco* ARE

In order to gain further insights into the mechanism of hormone-mediated de-repression, we also determined the structure of AvaR1 bound to a synthetic oligonucleotide derived from the autoregulator responsive element (ARE) sequence. Prior DNase foot-printing analysis established the identity of the ARE located upstream of the *aco* gene, however, this response element is pseudopalindromic.^14^ Crystallization efforts with the symmetric AvaR1 homodimer yielded crystals that did not diffract beyond 8 Å, presumably as a result of the asymmetry of the ARE operator. Efforts using an artificial palindromic sequence derived by inverting and repeating each half of the pseudo palindrome yielded crystals that diffracted to 3.09 Å resolution (Figure 3D, Figure S2, Table S1), and the structure was determined by molecular replacement. As a result of the use of this symmetric DNA, each homodimer in the crystallographic asymmetric unit is bound to a monomer from an adjacent ARE.

The structure shows that each DNA binding domain (DBD) interacts with one half of the palindrome of the DNA duplex. Numerous contacts are formed between helix α1 of the DBD and the duplex, including Arg5, which insert into the major groove and interact with Thy7, as well as between Lys43 and Ade15 of the ARE (Figure 3D). Additional non-specific interactions include those between Thr10, Thr42, and Tyr47 and the backbone phosphate of the duplex. A comparison of the ligand-binding sites with that in the hormone-bound structure reveals that the binding pockets is further occluded through movements of Trp127, as well as the loop harboring Gln64, consistent with the roles of these residues effecting conformational movements of the DBD upon binding of the hormone.

### Genome mining informs on putative butenolide regulatory biosynthetic pathways

Given the improved stability of butenolides over γ-butyrolactones, we speculate that these hormones may prove more amenable at attempts to activate antibiotic production. In order to identify actinobacterial strains that are under butenolide regulatory control, we sought to use a bioinformatics approach based on identification of the corresponding receptor. However, as shown in Figure 3A, the sequence similarity between bona fide γ-butyrolactone receptors like ArpA, and butenolide receptors is high (40% sequence identity with AvaR1) precluding such analysis. Orphan receptors called pseudo γ-butyrolactone receptors that are activated by multiple ligands likewise share between 40% sequence identity with AvaR1, confounding simple sequence-based analysis. Prior efforts to discriminate between receptor classes have not proven to be fruitful, and phylogenetic analyses failed to discriminate between positive and negative regulators.

In an effort to identify other actinobacteria that are under butenolide regulatory control, we utilized synteny of the putative butenolide biosynthetic genes to distinguish between receptor clades. We first used the Enzyme Similarity Tool (EST) from the Enzyme Function Initiative to create a Sequence Similarity Network (SSN) of all members of the receptor class. Using an E-value cutoff of 10^−70^ produce an SNN in which characterized receptors of the various GBL families were segregated with mutually exclusive co-localization (Figure 4A). We used the resultant SSN as input for the EFI Genome Neighborhood Network (GNN) tool to identify nodes that are co-localized next to genes with PFams that are associated with putative avenolide biosynthetic genes. Knowledge that the butenolide receptors often regulate the production of its own ligand further enabled this approach and allowed for inferences into the class of ligands produced and recognized by receptors that have yet to be characterized. Because the EFI tools are only integrated with the Uniprot database, we also manually scoured through sequences in Genbank for similar operonic architecture to identify putative butenolide receptors based on genomic context.

**Fig. 4.**
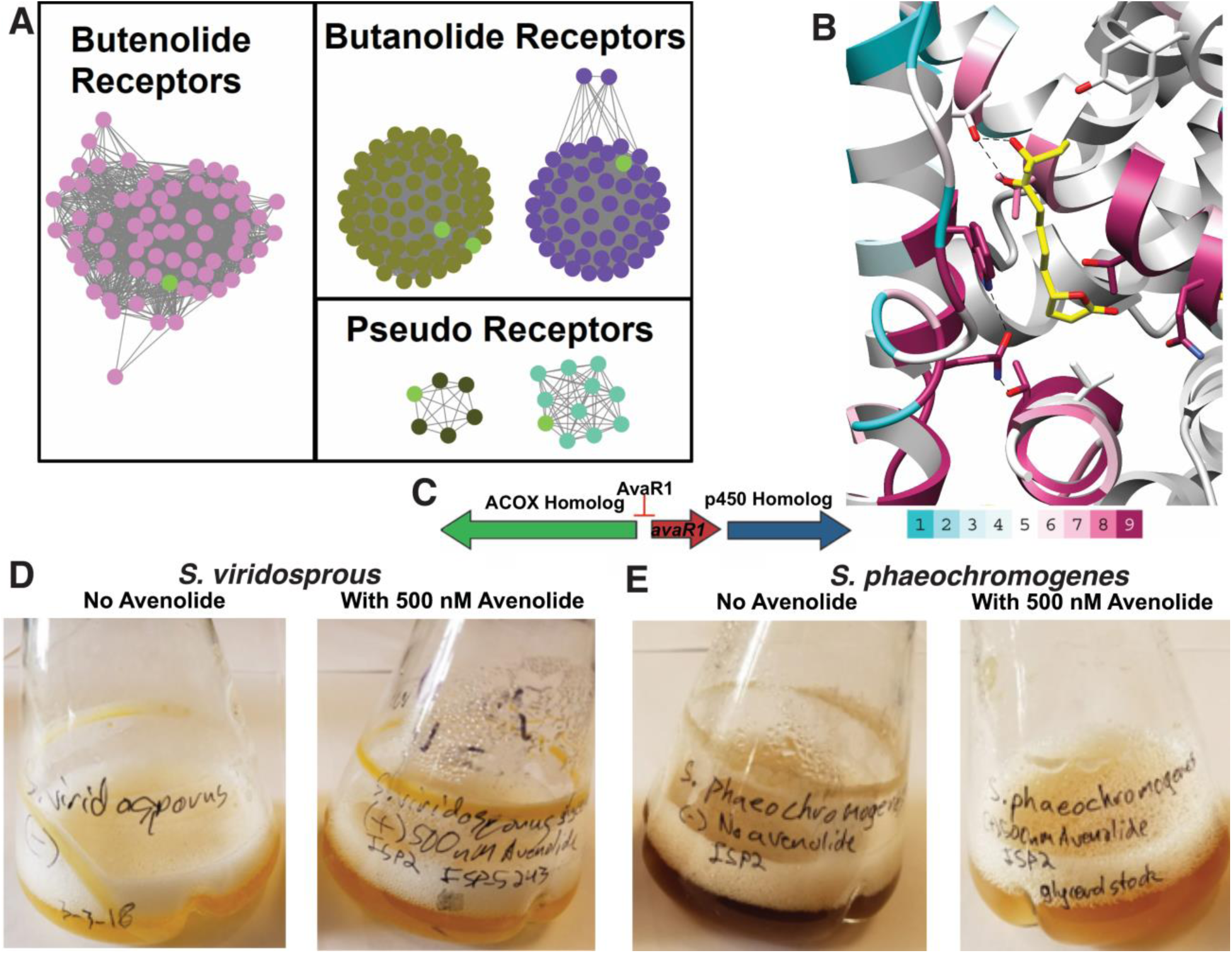
Sequence Similarity Networks of homologs gene clusters. **A**. SSN showing the relationship between different clades of putative butenolide receptors. Blue and purple (left box) are butanolide receptors; pink (middle box) are identified butanolide receptors and gold and light pink (right box) are the pseudo receptors. Characterized receptors are shown in yellow. **B.** Conservation of sequences amongst 90 putative butenolide receptors identified by bioinformatics mapped onto the structure of AvaR1. The color range indicates least conserved (cyan) through highest conserved (purple). **C**. Genomic synteny used to cull sequences for the SSN. **D** and **E**) Effects of avenolide on two actinobacterial strains identified to contain butenolide receptors based on the SSN analysis.

The aforementioned analysis yielded a set of 90 putative actinobacterial genomes (Table S2) that harbor a putative butenolide receptor likely under butenolide regulatory control (Figure 4A). We emphasize that, while this is orders greater than the currently known number of butenolide receptors, this number represents a significant underestimate and the actual number of receptors is well above this number but are not identified given the limitations of our approach. Mapping of the sequence conservation amongst these 90 receptors onto the cocrystal structure of hormone bound to AvaR1 reveals residues Gln64, Thr108, and Trp127 that is proposed to couple ligand binding to domain shift are conserved among all of these sequences (Figure 4B). The conservation score is highest at residues involved in interactions with the lactone, but those that interact with the alkyl tail are more divergent. These data are consistent with the observations that the lactone ring and C10 hydroxyl are general features of butenolides, whereas the length and branching of the alkyl tail vary significantly.

As a cursory validation of our bioinformatics results, we tested two of the actinobacterial strains identified as regulated by butenolides for phenotypic changes when the hormone is added to the growth cultures. Both *Streptomyces viridosporus* and *Streptomyces phaeochromogenes* contain a biosynthetic operon that harbor genes homologous to those purported to be involved in butenolide biosynthesis, located adjacent to an AvaR1 homolog (Figure 4C). Notably, growth of these strains in the presence and in the absence of added avenolide resulted in a change in the color of the culture medium. The change in color may be due to the production of new secondary metabolites and/or chromophores (Figure 4D, E). Given that the cognate hormone for each of these strains is likely not avenolide but rather a derivative, the changes in culture media in the presence of the hormone is consistent with the role of a butenolide receptor regulating secondary metabolite production in these strains.

## CONCLUSION

Although the bacterial hormone A-factor was discovered nearly a half century ago, significant gaps remain in our understanding of how these signaling molecules regulate gene expression. The discovery of the regulation of secondary metabolite biosynthesis by γ-butyrolactones pushed efforts to use these molecules as *ex vivo* effectors to induce otherwise silent biosynthetic gene clusters but with little success. Presumably, the labile nature of the lactone ring, as well as the often pleiotropic effects that γ-butyrolactones induce have subverted efforts to utilize these small molecules as chemical inducers. In contrast, the structurally related butenolides show improved stability under strongly acidic and basic conditions. However, utility of butenolides in biotechnology efforts is limited by the inability to access these molecules.

Here, we present here an efficient and convergent 22 step total synthetic route for the production of avenolide, which can be extrapolated for the total synthesis of other members of the butenolide class of small signaling molecules. This effort allowed for detailed structure-function studies of the corresponding hormone receptor, including the first crystal structure of any GBL type receptor bound to its cognate ligand. The structural data informs on the mechanism by which hormone binding induces a conformational change in the AvaR1 receptor, resulting in the formations of a dimeric assembly that occlude efficient DNA binding. We also elaborate a bioinformatics strategy using the genomic neighborhood context of butenolide biosynthetic genes as a marker to identify 90 actinobacterial strains that are likely under the regulatory control of butenolides. Addition of avenolide to the growth media for two representative strains results in changes in the color of the culture supernatant. These results support the validity of our bioinformatics approaches and set the framework for further efforts towards the use of butenolides to active antibiotic biosynthesis in otherwise silent gene clusters.

## Acknowledgements

We thank Keith Brister and colleagues at LS-CAT (Argonne National Labs) for facilitating X-ray data collection.

## MATERIALS AND METHODS

Total synthetic schemes, experimental procedures, and validation of relevant synthetic intermediates are provided in the Supplemental Data.

### Expression, purification, crystallization and structure determination of AvaR1

Wild-type protein AvaR1 was amplified from *Streptomyces avermitilis* genomic DNA by PCR using primers based on the published sequence of the polypeptide and inserted into a ligation independent cloning vector for expression in *E. coli* as a maltose binding protein (MBP)-tagged fusion. The resultant plasmid was transformed into *E. coli* containing the Rosetta plasmid for protein expression. AvaR1 was produced by growing the cells in shaking flask of LB media at a temperature of 37 °C. When the cells reached an O.D._600_ of 0.6, the cells were cooled on ice for 15 minutes. Following the addition of 0.5 mM IPTG, the cells were placed in an 18 °C shaking incubator for 18 hours. The cells were then harvested by centrifugation and resuspended in a buffer composed of 500 mM NaCl, 20 mM Tris base (pH 8.0), and 10% glycerol. Resuspended cells were lysed by homogenization and the lysate was centrifuged at 14,000 rpm to remove cell debris. The cleared cell lysate was loaded onto a HisTrap column, which was subsequently washed with 1 M NaCl, 30 mM imidazole, and 20 mM Tris base (pH 8.0). MBP tagged AvaR1 was eluted using a linear gradient beginning with 1 M NaCl, 20 mM Tris base (pH 8.0), and 30 mM imidazole and ending with 250 mM imidazole. Pure fractions, as judged by SDS-PAGE, were combined and diluted two-fold before treatment with thrombin (final ratio of 1:100 (w/w) for 18 hours at 4 °C to cleave the N-terminal tag. Tag-free AvaR1 was concentrated and loaded onto a size exclusion column (Superdex S75 16/60) pre-equilibrated with containing 100 mM KCl and 20 mM HEPES free acid (pH 7.5). Pure fractions were collected and concentrated to 25 mg/mL before storage in liquid nitrogen. Production of SeMet labeled AvaR1 was carried out by repression of methionine synthesis in defined media supplemented with selenomethionine.^27^

Preliminary crystals of AvaR1 were obtained using a sparse matrix screen. Diffraction quality crystals were grown using hanging drop vapor diffusion with 13.5 mg/mL AvaR1 was added to mother liquor containing 12% PEG 1000, 0.1 M sodium citrate tribasic dehydrate (pH 4.2), 0.2 M LiSO_4_ and 4% (v/v) tert-butanol in a 1:1 ratio at incubated against the same solution at 4 °C. Crystals were improved through multiple rounds of micro-seeding. The SeMet-AvaR1 crystals were obtained using 12 mg/mL protein added to a 1:1 ratio of 30% PEG MME 2000, and 0.15 M KBr. Crystals were vitrified by direct immersion without the addition of any cryo-protectives.

All diffraction data were collected at Argonne National Laboratory (IL). The autoPROC^28^ software package was utilized for indexing and scaling of the diffraction data. Initial phases for AvaR1 were obtained using anomalous diffraction data collected on crystals of SeMet labeled protein. Initial models were built using Phenix and Parrot/Buccaneer. Manual refinements were completed by the iterative use of COOT^29^ and Phenix.refine. Cross-validation was utilized throughout model building process in order to monitor building bias. The stereochemistry of all of the models was routinely monitored using PROCHECK. Crystallographic statistics are provided in Table S3 of supplementary information.

For co-crystallization of the hormone bound complex, purified AvaR1 (14 mg/ml) was incubated with 3 mM avenolide for 30 minutes on ice. Co-crystals were obtained by vapor diffusion methods and initial crystals were obtained in Index D5 (25% PEG3350 and 0.1M sodium acetate trihydrate pH 4.5). Well diffracting crystals were produced through optimization to a final solution of 23% PEG3350 and 0.1 M sodium acetate trihydrate pH 4.5 at 4 °C using hanging drop crystallization. Crystals were submerged briefly in the crystallization media supplemented with 25% ethylene glycol prior to vitrification in liquid nitrogen. The coordinates of apo AvaR1 were used to determine crystallographic phases.

Co-crystallization of AvaR1 with different oligonucleotide sequences designed based on the AvaR1 DNA binding site upstream of *aco* gene. Purified and concentrated dimeric protein (14 mg/mL) was incubated with individual oligonucleotide duplexes (Supporting Table S1) in 1:1.2 molar ratio, for 30 minutes on ice. Palindromic DNA sequences were first self-annealed and then double stranded DNA was used from 1 mM stock prepared in 20 mM MgCl_2_, 50 mM Tris pH 8.0 buffer. Buffer, DNA and protein were added in respective order. For some oligonucleotides, white turbid solution was obtained as soon as protein was added, addition of a few microliters of ammonium acetate and incubating at either room temperature or on ice, produced clear solution. Crystallization trays were set up at 4 °C and every oligonucleotide crystallized in different conditions (Figure S2B). Ethylene glycol (25% v/v) was used as cryoprotectant prior to vitrification of crystals for all AvaR1-oligonucleotide co-crystals. The oligonucleotide sequences used are listed in Table S1 and the sequence that produced diffraction quality crystals have been shown in Figure S2 of supporting information.

### Identification of Putative Butenolide Biosynthetic Clusters

Using the AvaR1 amino acid sequence as a handle, tools from the Enzyme Function Initiative (EFI) were used to first create a Sequence Similarity Network (SSN) of 10,000 Uniprot sequences. Once an SSN was created, an iterative process was undergone to find an E-value to ensure characterized receptors of the various GBL families of receptors result with mutually exclusive co-localization. The resulting E-value was 10^−70^. Using this SSN, the data was run through the EFI’s Genome Neighborhood Network (GNN) webtool. Network visualization was performed in Cytoscape.^30^ Using Pfams associated with putative avenolide biosynthetic genes along with the knowledge that this family of receptors often regulates their own ligand production, inferences are made as to the class of ligand produced and recognized by uncharacterized receptors. Because the EFI webtools are only integrated with the uniprot database, we also manually scoured through a number of Genbank sequence results derived from BLAST analysis for proper genomic context relative to butenolide production. These BLAST searches were using the sequences of any putative butenolide biosynthetic genes as handles. What was considered proper genomic context necessary for butenolide biosynthesis was a TetR_N Pfam receptor surrounded by a gene in the p450 Pfam (PF00067), and either an Acyl-CoA_dh_1 (PF00441) Pfam gene, an Acyl-CoA-dh_2 (PF08028) Pfam gene, or an ACOX (PF01756) Pfam gene. List of these 90 homologous strains is provided in Table S2 of supporting information.

### Isothermal Titration Calorimetry

ITC measurements were performed at 25 °C on a MicroCal VP-ITC calorimeter. A typical experiment consisted of titrating 7 µL of a ligand solution (80 µM) from a 250 µL syringe (stirred at 300 rpm) into a sample cell containing 1.8 mL of AvaR1 solution (8 µM) with a total of 35 injections (2 µL for the first injection and 7 µL for the remaining injections). The initial delay prior to the first injection was 60 s, with reference power 10 μCal/s. The duration of each injection was 16 s and the delay between injections was 400 s. All experiments were performed in triplicates. Data analysis was carried out with Origin 5.0 software. Binding parameters, such as the dissociation constant (K_d_), enthalpy change (∆H), and entropy change (∆S), were determined by fitting the experimental binding isotherms with appropriate models (one-site binding model). The ligand stock solution was prepared at 10 mMThe buffer solutions for ITC experiments contained 300 mM KCl and 20 mM HEPES pH 7.5.

### Total Synthesis of Avenolide

The experimental procedures were adopted from the publication by Uchida et.al.^20^ with further optimizations and modifications that have been stated. Detailed experimental procedures for relevant intermediates are specified in materials and methods section of supplementary information.

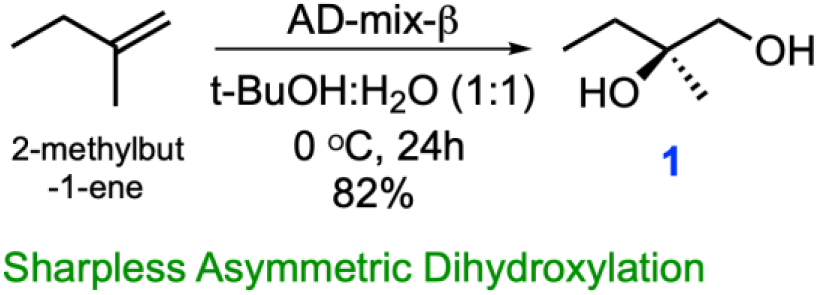

*(R)-2-Methylbutane-1,2-diol* **(1):** To a stirred solution of the 2-methyl-1-butene (3.75 g, 53.47 mmol) in t-BuOH: H_2_O (1:1, 400 ml) were added K_3_Fe(CN)_6_ (52.81 g, 160.41 mmol), K_2_CO_3_ (22.17 g, 160.41 mmol), K_2_OsO_4_(OH)_4_ (197 mg, 0.54 mmol, 1 mol%) and (DHQD)_2_PHAL (416.5 mg, 0.54 mmol, 1 mol%) at 0 °C under Ar atmosphere. The reaction mixture was stirred for 24 h at 0 °C using Ar balloon. The reaction was quenched with a saturated aqueous solution of Na_2_S_2_O_3_ and the aqueous phase was extracted with EtOAc (2×1 L). The water layer was thoroughly washed with EtOAc and combined organic extracts were washed with brine (saturated NaCl) and dried over anhydrous Na_2_SO_4_ and concentrated in vacuo. The residue was purified by flash column chromatography with a gradient from 30% EtOAc/hexanes to 70% EtOAc/hexanes to 10% MeOH/DCM, to afford **1** (3.5 g, 82%) as a colorless oil. **1** was obtained in single step from commercially available 2-methylbut-1-ene using Sharpless asymmetric dihydroxylation^21^. Spectroscopic characterization parameters agreed with the reported mass and chemical shift values.

[α]^25^ +4.96 (c 1.0, CHCl_3_); 1HNMR (500 MHz, CDCl_3_) δ3.47 (brs, 2H), 3.43 (d, *J* = 11.1 Hz, 1H), 3.37 (d, *J* =11.1 Hz, 1H), 1.51 (q, *J* =7.3 Hz, 2H), 1.10 (s, 3H), 0.89 (t, *J* =7.6 Hz, 3H); 13C-NMR (125 MHz, CDCl_3_) δ73.6, 69.3, 31.2, 22.5, 8.2 HRMS (ESI+, TFA-Na) calcd for C_5_H_12_NaO_2_ 127.0735 [M+Na]^+^, found m/z 127.0740.

For synthesis of aldehyde fragment **12**, the PMB protected intermediates were synthesized following protocol reported by Uchida *et.al.* Iodo alkene fragment **11** was also synthesized following the reported protocol by substituting the use of TBS protecting group with TBDPS to enable easy monitoring of reaction by TLC visualization under UV light. Epoxy fragment **10** was synthesized through intermediates **14**,**15** and **16**, as described in the supporting information.

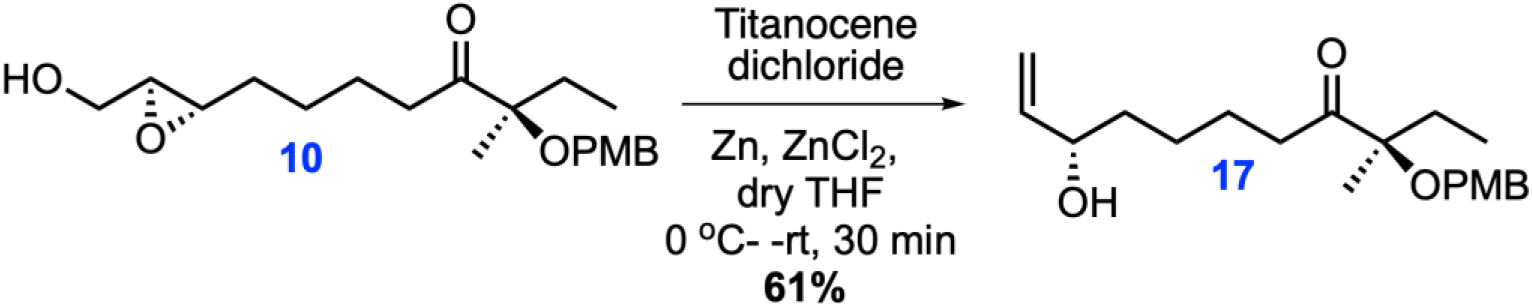

*(R)-8-((2S,3S)-3-(Hydroxymethyl)oxiran-2-yl)-3-((4-methoxybenzyl)oxy)-3-methyloctan-4-one* (**17**): Following the protocol from the reported literature^22^ anhydrous ZnCl_2_ (2 mL, 1 M in Et_2_O, 2 mmol) and zinc powder (350 mg, 6.72 mmol) were added to a red solution of Cp_2_TiCl_2_ (1.26 g, 5.05 mmol) in anhydrous THF (15 mL). The solution was stirred for 1h at room temperature until it turned green. Epoxide **10** (590 mg, 1.68 mmol) in anhydrous THF (5 mL) was then added to the resultant mixture. After stirring for 30 min at rt, the reaction was quenched with aqueous HCl (1.0 M, 3 mL) and the mixture was extracted three times with Et_2_O (4 mL). Collected Et_2_O fractions were combined and washed with water, 10% aq. NaHCO_3_, water and brine, dried over Na_2_SO_4_ and filtered and concentrated under reduced pressure. Obtained residue was purified using flash column chromatography on silica gel (25% EtOAc/hexane to 40% EtOAc/hexane) to obtain pure allyl alcohol compound **17**. ^1^H-NMR (500MHz, CDCl_3_) δ7.27 (d, *J =* 8.8Hz, 2H), 6.89 (d, *J =* 8.8Hz, 2H), 5.87-5.81 (m, 1H), 5.21 (ddd, *J=*17.2, 1.4, 1H), 5.10 (ddd, *J=*10.4, 1.4, 1H), 4.33 (d, *J =* 10.8Hz, 1H), 4.29 (d, *J =* 10.8Hz, 1H), 4.14-4.06 (m, 1H), 3.81 (s, 3H), 2.66 (dt, *J =* 7.3, 4.3Hz, 2H), 1.84–1.70 (m, 2H), 1.59–1.34 (m, 6H), 1.33 (s, 3H), 0.84 (t, *J =* 7.5Hz, 3H); ^13^C-NMR (500MHz, CDCl_3_) δ215.2, 159.0, 141.3, 128.9, 128.9, 114.9, 114.0, 113.8, 84.8, 73.2, 65.3, 55.5, 37.1, 36.8, 29.4, 25.3, 23.5, 20.2, 8.1; HRMS (ESI+, TFA-Na) calcd for C_20_H_30_NaO_4_ 357.2042 [M+Na]+, found m/z 373.2032.

This resulted in a simplified protocol for the synthesis of allyl alcohol **17** which otherwise was reported to be made in 2 additional reaction steps starting from epoxy **10** from. Acrylic group was added to ally alcohol **17** using DDQ by the reported procedure and followed by ring closing metathesis reaction to yield stereospecific (4*S*, 10*R*)-avenolide, **13.** Detailed synthetic schemes, experimental procedures and yields have been reported in materials and methods section of supporting information. Obtained ^1^H and ^13^C NMR data supports the reported values, thus the spectra are provided only for key intermediates.

